# Influence of spatial structure on protein damage susceptibility – A bioinformatics approach

**DOI:** 10.1101/2020.03.03.973099

**Authors:** Maximilian Fichtner, Stefan Schuster, Heiko Stark

**Affiliations:** Matthias Schleiden Institute, Department of Bioinformatics, Friedrich Schiller University Jena, Ernst-Abbe-Platz 2, 07743 Jena, Germany; Institute of Zoology and Evolutionary Research, Friedrich Schiller University Jena, Erbertstraße 1, 07743 Jena, Germany

**Keywords:** Amino acid score, 3-dimensional protein structure, protein shell, protein core, protein damage, protein aging, concave hull

## Abstract

Aging research is a very popular field of research in which the gradual transformation of functional states into dysfunctional states are studied. Here we only consider the molecular level, which can also have effects on the macroscopic level. It is known that the proteinogenic amino acids differ in their modification susceptibilities and this can affect the function of proteins. For this it is important to know the distribution of amino acids between the protein surface/shell and the core. This was investigated in this study for all known structural data of peptides and proteins. As a result it is shown that the surface contains less susceptible amino acids than the core with the exception of thermophilic organisms. Furthermore, proteins could be classified according to their susceptibility. This can then be used in applications such as phylogeny, aging research, molecular medicine and synthetic biology.

## Introduction

Aging research is a very popular field of research that focuses on macroscopic and microscopic alterations during aging. Aging is a biological process in which a functional state is gradually transformed into a dysfunctional state[1–3]. Macroscopic changes can be skin aging, reduced mobility and organ damage (heart failure, autoimmune diseases such as age-related macular degeneration, neurodegenerative diseases such as Alzheimer’s disease). At the cellular (microscopic) level, the changes affect the signalling and metabolic pathways as well as many larger molecules. The changes of the molecules are of decisive importance as they can accumulate in an organism and lead to macroscopic changes (e.g. lipofuscin, age pigment in the skin;[4]). For many molecules there are already detailed studies available. Known are for misfolded proteins caused by mutation: α-synuclein, cystic fibrosis transmembrane conductance regulator[5], peripheral myelin protein 22[6], huntingtin (Htt)[7], ataxin-3, Down syndrome critical region 1[8]. There are fewer investigations for misfoldings due to non-enzymatic modifications. Nevertheless, it is worth mentioning that ageing is characterized not only by a decline in function but also by a remarkable robustness of many features such as the hematocrit value [9], body temperature, overall immune memory etc.

### Score based estimation of peptide and protein susceptibilities

In our previous paper [10] we assembled score tables and proposed an approach to quantify peptide and protein susceptibility. For the score tables we have selected known protein modifications from four literature sources[11–14]. On this basis the susceptibilities for the 20 standard amino acids (AAs) were determined. In a second step, the susceptibilities were weighted by text mining. In a third step we weighted only with text mining, without the consideration of the modification table. Finally, an average of all scores was calculated.

At first this was applied without consideration of the three-dimensional (3D) structure. Merely a distinction between peptides and proteins was made using a threshold of 100 AAs. It can be expected that peptides have no core in their spatial structures due to their short length. For simplification, only if necessary we make a distinction between peptides and proteins, otherwise we just use the term proteins for both in the following. In the present paper, we take the 3D structure fully into account.

A similar approach in terms of structure scoring that connects well with our research has recently been published. There, the authors score the structures of proteins for their susceptibility to aggregate and call it AggScore[15]. An additional study (review) addressed AAs in the context of oxidation, which also ranked highest in our score (cysteine, tyrosine and tryptophan)[16]. They also mention the problem in connection with storage of biotherapeutics for longer durations. It is notable that some other AAs mentioned in [16] like histidine, methionine and phenylalanine were just ranked average in our score.

### Spatial protein structures

While we mainly consider the primary structure (complete amino acid chain) in the previous paper[10], here we take spatial information into account. Obviously, the susceptibility of an AA to spontaneous modification depends on its localization within the protein. AAs in the protein shell (further called protein surface) are much more easily accessible to, for example, reactive oxygen species, than those in the core. Since the secondary and super-secondary structures transition relative quickly into the tertiary structure they seem to be, In terms of the susceptibility, only relevant for peptides or small proteins where they are fully accessible. In Table 1 we give a short view on the secondary structures alone, that means without consideration of their spatial arrangement in the protein[17–20].

**Table 1:**
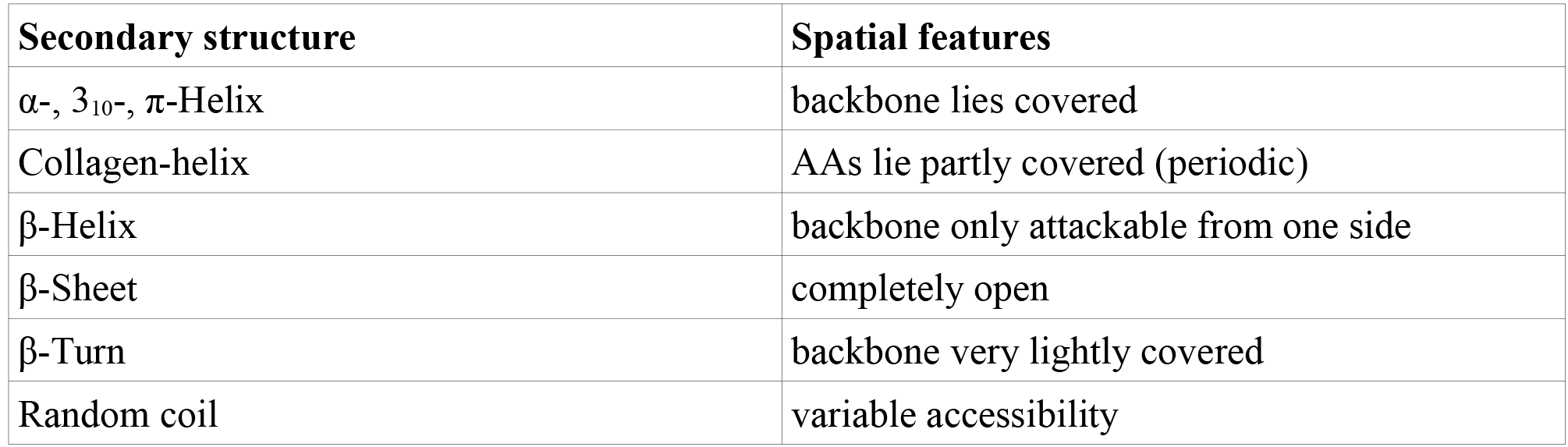
Summary of secondary structures and possible spatial susceptibility. The spatial features are derived from the chemical structures.

Within the tertiary structure the composition of secondary structures is decisive for the accessibility of the AAs. If, for example, there are more α-helices outside than β-sheet structures, a different impact on the surface is to be expected. In the calculation of the whole protein surface this issue is already addressed.

Furthermore, the surface can be covered by other structures. In the quaternary structure, individual proteins form protein complexes. After the formation of a protein complex, a new protein surface is formed. In addition, a distinction can be made between the complete surface and the accessible surface. For example, proteins can be embedded in membranes and are thus only partially vulnerable to the respective micro-environments (e.g. mitochondria, chloroplasts, …). Especially for enzymes, susceptible/functional areas can also be hidden in pockets. However, here we focus only on the tertiary and possibly also the quaternary structure, since this represents the final form of the proteins. It should be noted that the spatial structure is subject to fluctuations and this can lead to differently measured data[21].

### 3D approach

The idea was to make a 3D approach where only the AAs which are lying on the outside of the protein surface are considered for the calculation. There are a number of algorithms for the calculation of protein geometries, which calculate the protein surface, volumes and pockets[22,23,21,24,25]. Here the outer AAs need to be identified. In contrast, the remaining AAs form the protein core. We assume that more susceptible AAs in the protein core are protected by the AA on the protein surface and are therefore more susceptible to modification. Thus, the protein surface AAs should be less susceptible to modification. It is also known that certain AAs protect proteins [26–28].

Additionally one can make a distinction between the backbone and the side chains. Depending on the folding, the respective parts are accessible from the outside. The same applies to pockets, which, depending on size and depth, offer more vulnerale surface.

### Hypothesis

Compared to Fichtner et al. [10], it can be expected that there will be a difference between the whole and parts of the protein. The AAs in the core are protected by the protein surface and could have more susceptible AAs. On closer inspection, the surface can also be analysed with regard to its differences with and without backbone and with and without side chains. It is to be expected that specific protein families form clusters with regard to their susceptibility.

## Methods

### Preprocessing

As a basis for the analysis, the spatial structures of all molecules (139,291) deposited in the Protein Data Bank (PDB) were downloaded (see Suppl. 1). In the preprocessing only the compositions which contain AAs in their sequence were considered (Suppl. 1). In addition, the data sets may include also ligands and water. These were retained for the calculation, as they form the outermost surface and can act as protection. A direct comparison for multiple spatial structures is too combinatorial challenging (comparing all atoms with each other), that is why only one spatial structure per entry was considered (Suppl. 1). Additionally, the entries with redundant sequences may differ in their spatial structure, resulting in different surface and core compositions, and were therefore not summarized here. For our analysis we concentrate only on the PDB data. A connection with further databases would be possible. For example to investigate the influence of domains.

### Whole protein

We have developed nine different approaches to compare the influence of the spatial structure. The simplest one, ‘whole protein’ (WP) considers all AAs (like the theoretical approach [10]). Based on these results, we again calculated the susceptibilities for all proteins for which we had the spatial data (Suppl. 1). For that purpose we only used the score ALL of Fichtner et al. [10]. This allows us to compare parts (surface, core) with the whole protein in terms of susceptibility. Score ALL is a mean of six scores with different focuses in terms of weighting and data base. With the mean it was tried to combine the best characteristics of the scores.

### Protein core and protein surface

By defining the protein parts, we can mainly examine the ‘protein core’ (PC) on the one hand and the ‘protein surface’ (PS) on the other. In addition, we make a distinction in the surface between and ‘surface side chains’ (SSC) and ‘surface backbone’ (SBB) which only consists of the backbone elements in the surface. Depending on the spatial structure the corresponding protein parts can lie outside or inside. For our scores we have only considered the side chains. In the protein surface, the backbone may or may not point outwards and be susceptible. That is why we have devised another approach where we analyse the whole surface and then we exclude the backbones from the analysis (SSC). For the sake of completeness, we have also devised an approach in which the side chains are subsequently removed (SBB). Thus, it is possible to analyse the number of side chains of individual proteins in comparison to the backbone on the protein surface.

For the determination of the PS, SSC, SBB the concept of the ‘concave hull’ is used. This is a complex problem in information theory. That is because there is no agreed definition for it, since we have to decide at which point we stop chasing deeper gaps. That is why the concave protein surface will be defined depending on the requirements of the problem in question. For this purpose we chose a small radical (hydroxyl radical), which is reactive at any pH value. With the help of the software ‘Avogadro’ Version 1.1.1[29,30] we determined the minimum distance for chasing deeper gaps. While the Van-der-Waals radii of bound proteinogenic atoms are larger, the radii of isolated atoms are between 1.1 Å and 1.8 Å [31,32]. Here an oxygen radical (1.52 Å) was moved through two carbon atoms (1.7 Å) and the distance was measured (6 Å to 7 Å). These values are to be understood as the lowest limit. It should be noted that this is not a fixed limit and is therefore only an approximation. A rough formula could be: X + 2 * ½ Y = X + Y where X is the Van-der-Waals diameter of the penetrating atom and Y is the Van-der-Waals diameter of the two surface atoms that are not bound with each other. The respective formula for radii would then be 2 X + 2 Y.

For the definition of our PS (biological term) we use the standard Graham Scan algorithm[33]. The Graham Scan is an algorithm for calculating the convex hull (mathematical term) of a finite set of points. We calculate the convex hull for one protein based on all atoms of the protein. After that we divide each edge by the minimal distance (6 Å to 7 Å) and the new nodes to the closest points are calculated. In this way, we obtain the concave hull, but only for topologies with genus zero (i.e. without holes). The surface contains the external atoms of the molecule and in a second step the corresponding AA to these atoms were determined (Fig. 1). After this assignment, surface and core of the protein can be separated. The algorithm described here was implemented in the program Cloud2 Version 14.3.20 (Heiko Stark, Jena, Germany, URL: https://starkrats.de) for the differentiation of the different protein parts (Suppl. 1).

**Figure 1:**
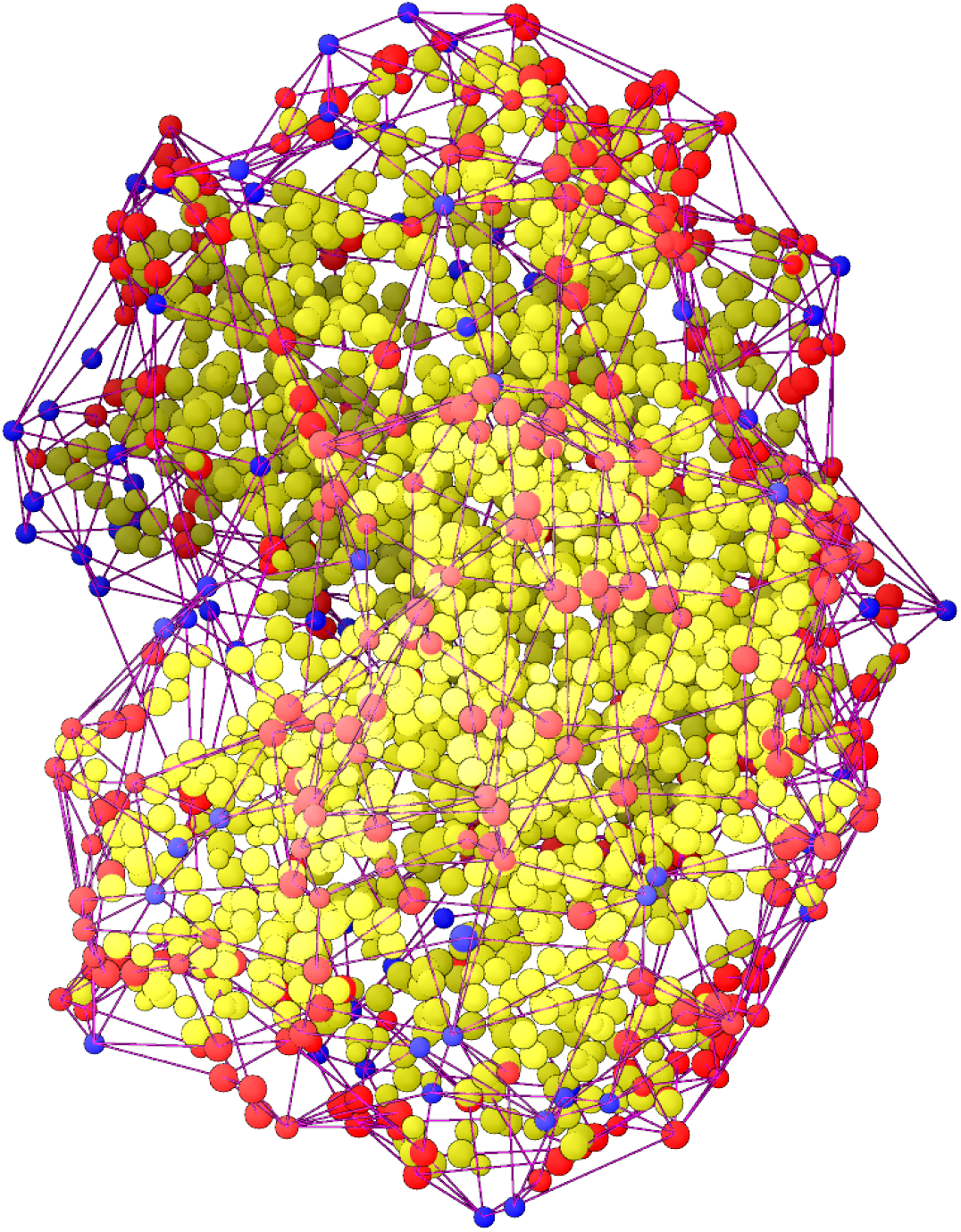
Representation of the amino acids and water components of the protein oxyhaemoglobin (PDB entry: pdb1hho). A dot represents the averaged centre of an amino acid (red when on the protein surface (PS), yellow in the protein core (PC)) or water molecule (blue). Red lines show the cross-linking by the surface calculation, by which the surface (concave hull) is defined. All unlinked dots are assigned to the PC.

### Scoring

As a first step, the entries with multiple conformations were removed for reasons of complexity. This included 8039 (5.77%) entries. From this data, an AA count was performed for validation. In addition, the sequences for the different protein parts (WP, PS, SBB, SSC, PC) were weighted with the score ALL of Fichtner et al. and a modified scoring program (Suppl. 1; [10]). In a second step the AA sequences with more than 5% X were removed (named with ‘X correction’; this makes 5.40 % of all entries). The 5% were taken from Fichtner et al. [10].

### Classification

Based on the annotations in the protein lists, a classification of special protein groups (collagen, cytochromes, ribosomes, …), organisms and enzymes was possible. This has been realized with the tool Enzyme2 Version 8.3.20 (Heiko Stark, Jena, Germany, URL: https://starkrats.de). For this purpose, the complete list was searched for specific patterns (e.g. enzymes – lyases, isomerases,… see Suppl. 5; organisms – human, mouse,… see Suppl. 6) and plots were created.

## Results

### Raw data

The first step was to evaluate the raw data for the various analyses (Tab. 2). It should be noted that WP contains all proteins. The calculation for PS resulted in as many entries as WP (since all proteins have a surface). The file size for 6 Å is larger than for 7 Å, since the penetration depth of the concave hull is larger, more AAs are found here (this applies to all calculated surfaces). With SBB almost no proteins are lost compared to PS. However, when comparing the file size of SBB with PS it is noticable that less AAs are selected. With SSC the number of proteins is reduced, i.e. there are proteins without an external side chain atom. In the 7 Å variant, this is even more pronounced because here the net has a lower density. When comparing the file size of SBB with SSC it is noticable that there are many more side chains outside than backbones. Not every protein has a core (PC), that is why here the number of proteins is lower compared to WP.

**Table 2:** Results of the number of proteins per calculation and file size. File sizes where taken without headers.

| Peptides/Proteins | Cavity 6 Å – count<br>[file size without header] | Cavity 7 Å – count<br>[file size without header] |
| --- | --- | --- |
| WP | 139,291<br>[111.5 MB] |  |
| PS | 139,291<br>[39.9 MB] | 139,291<br>[36.8 MB] |
| SSC | 138,921<br>[34.6 MB] | 138,921<br>[32.0 MB] |
| SBB | 139,256<br>[17.9 MB] | 139,243<br>[15.1 MB] |
| PC | 137,309<br>[71.8 MB] | 137,517<br>[74.9 MB] |

### Comparison linear versus spatial approach

The amount of molecules (139,291) is smaller compared to that analysed in Fichtner et al. [10] (422,091), since the spatial structure is not known for each molecule. A direct comparison of the histograms between these two studies shows a good correlation with regard to the normal distribution (Fig. 2). A further comparison of the distribution with respect to PS, SSC, SBB and PC shows that the normal distribution is shifting (Tab. 3 and 4). The SBB has the lowest mean and PC the highest. PS, SSC and WP are, in that order, situated in between. Our analysis showed that the choice of the AAs belonging to the PS is decisive for the way the susceptibility is calculated. There is a difference between 6 Å or 7 Å cavities (Tab. 3 and 4). The peptides show a higher standard deviation (SD) and lower mean values compared to the proteins.

**Figure 2:**
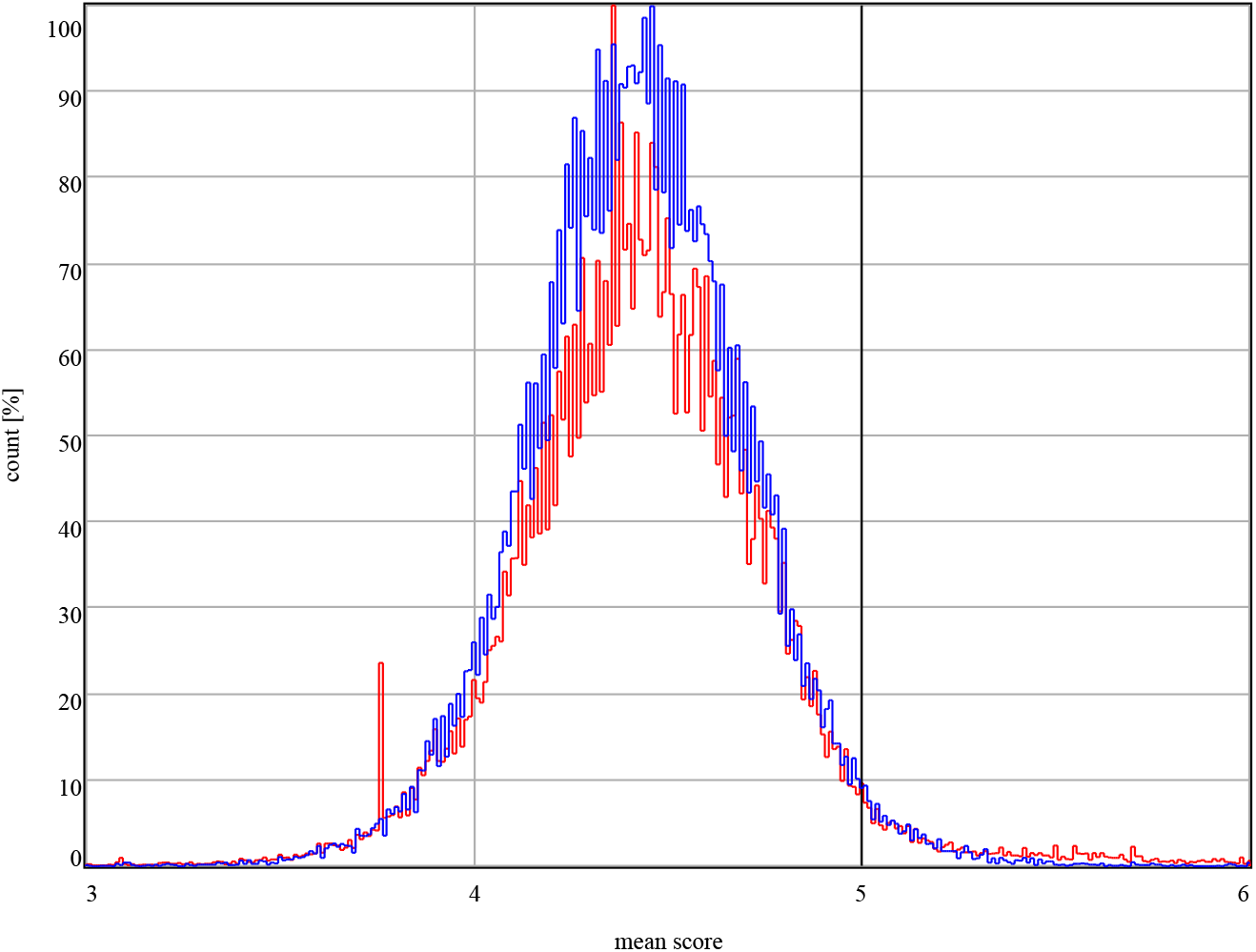
Comparison between the normalized histogram of [10] (blue) and the normalized histogram for the spatial approach (red). The red peak on the left (~3.757) is mainly due to multiple entries of endothiapepsin of the organism *Cryphonectria parasitica* in the data set. The second and highest peak (~4.357) is mainly due to lysozyme of the organism *Gallus gallus*. The black line indicates the mean score of a random artificial protein.

**Table 3:** Statistics of the influence of the spatial structure on the susceptibility for a cavity of 6 Å.

| Proteins<br>[Peptides] | Min | Median | Max | Mean | SD |
| --- | --- | --- | --- | --- | --- |
| WP | 0<br>[0] | 4.421<br>[3.910] | 8.594<br>[13.149] | 4.385<br>[2.710] | 0.498<br>[2.301] |
| PS | 0<br>[0] | 4.225<br>[4.193] | 8.403<br>[13.149] | 4.185<br>[3.780] | 0.555<br>[1.458] |
| SSC | 0<br>[0] | 4.261<br>[4.251] | 8.251<br>[14.750] | 4.228<br>[3.889] | 0.554<br>[1.379] |
| SBB | 0<br>[0] | 3.962<br>[3.921] | 6.194<br>[13.115] | 3.898<br>[3.815] | 0.626<br>[0.935] |
| PC | 0<br>[0] | 4.585<br>[4.625] | 8.582<br>[12.415] | 4.572<br>[4.522] | 0.465<br>[1.276] |

**Table 4:**
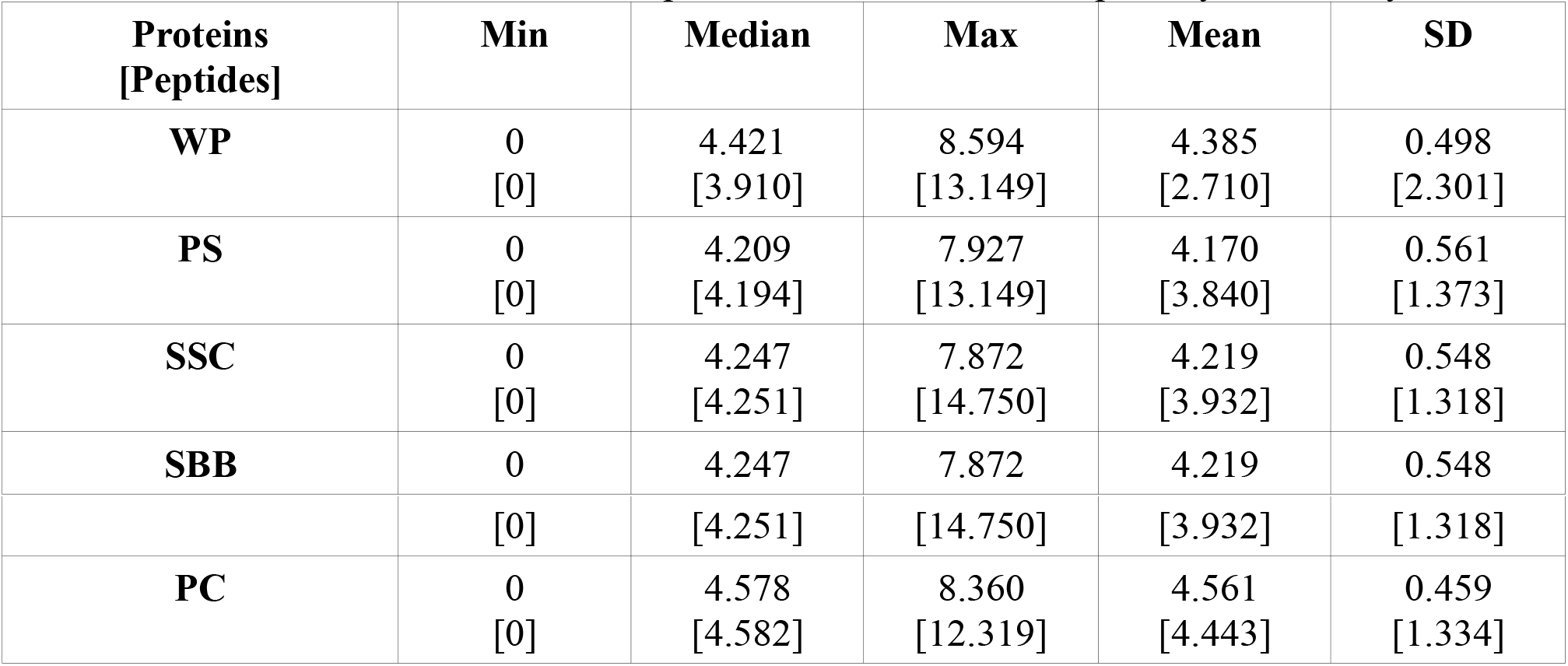
Statistics of the influence of the spatial structure on the susceptibility for a cavity of 7 Å.

| Proteins<br>[Peptides] | Min | Median | Max | Mean | SD |
| --- | --- | --- | --- | --- | --- |
| WP | 0<br>[0] | 4.421<br>[3.910] | 8.594<br>[13.149] | 4.385<br>[2.710] | 0.498<br>[2.301] |
| PS | 0<br>[0] | 4.209<br>[4.194] | 7.927<br>[13.149] | 4.170<br>[3.840] | 0.561<br>[1.373] |
| SSC | 0<br>[0] | 4.247<br>[4.251] | 7.872<br>[14.750] | 4.219<br>[3.932] | 0.548<br>[1.318] |
| SBB | 0 | 4.247 | 7.872 | 4.219 | 0.548 |
|  | [0] | [4.251] | [14.750] | [3.932] | [1.318] |
| **PC** | 0 | 4.578 | 8.360 | 4.561 | 0.459 |
|  | [0] | [4.582] | [12.319] | [4.443] | [1.334] |

### Validation of amino acid distribution in the protein surface

It is well known that, in cytosolic proteins, hydrophilic AAs are mainly found in the surface and hydrophobic AAs in the core[34]. These findings agree with our results and show that our algorithm correctly discriminates against the surface (Fig. 3; further analysis see Suppl. 2 and 3). Notice that in our data set the PC contains 64.3 % and the PS 35.7 % of all the AAs. The basic AAs lysine (K) and arginine (R) and the acidic AA glutamic acid (E) are hydrophilic and stand out due to a higher proportion in the PS. It should be mentioned here that K, R and E have a medium susceptibility, with E having the lowest value (Fichtner et al. [10]). Furthermore, the AAs glutamine (Q) and aspartic acid (D) show an equal distribution between PS and PC. In contrast to K, R and E, however, they show the lowest susceptibility. In addition, it is shown that the most susceptible AAs, i.e. tyrosine (Y), cysteine (C), tryptophan (W) and leucine (L), are represented above average in the core.

**Figure 3:**
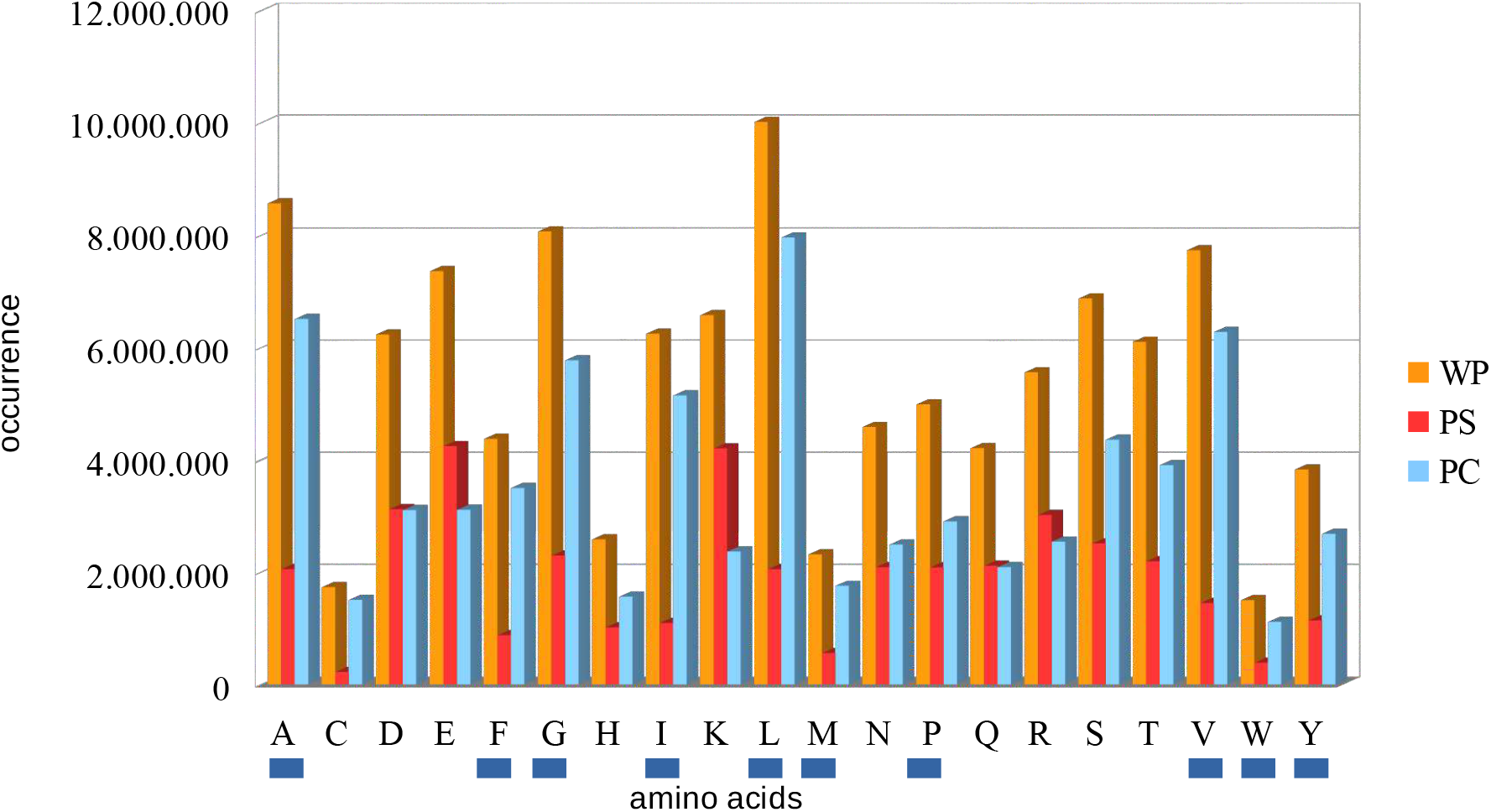
Real values of the occurrence of the 20 standard amino acids in the data and their distribution over protein surface (PS) and protein core (PC). The blue lines under the letters mark the hydrophobic amino acids. For further analysis see Suppl. 2 and 3.

### Comparison of protein core and protein surface

In the paper by Fichtner et al., reduced susceptibilities were shown for some proteins, among others for flagellin and spidroin[10]. A more precise differentiation between PS and PC shows that the surface of these proteins is less susceptible than the core and thus appears in the lowest susceptibility score for PS (Fig. 4; flagellin P value = 0.00471; spidroin P value = 0.243). For further analysis see Suppl. 4.

**Figure 4:**
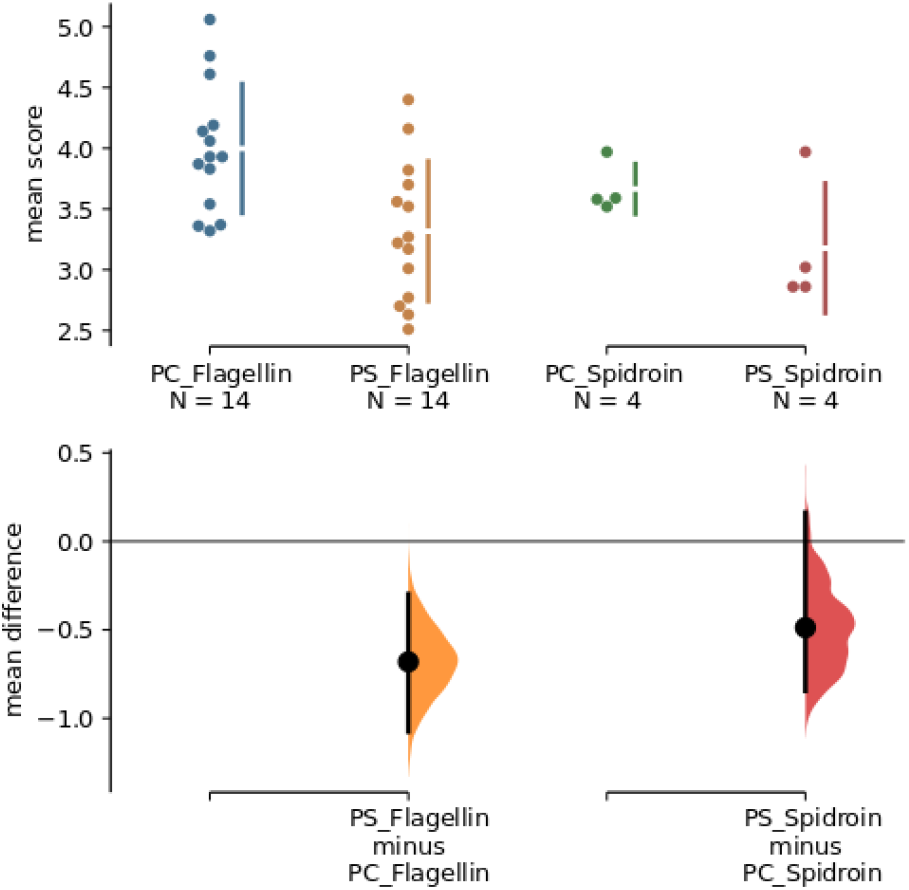
The mean difference of the protein core (PC) and protein surface (PS) comparisons for the flagellin and spidroin protein are shown in the above Cumming estimation plot. The raw data is plotted on the upper axe. Mean differences are depicted as dots; 95% confidence intervals are indicated by the ends of the vertical error bars. Each mean difference is plotted on the lower axes as a bootstrap sampling distribution (5000 bootstrap samples; confidence interval is bias-corrected and accelerated) [35]

When analyzing the surface, the location of the backbone or side chain is important for the analysis. For example, the heatshock protein shows that the proteins significantly influence the susceptibility calculation for the backbone (Fig. 5; SBB-PS P value 7.1e-08; SCC-PS P value 0.327). However, this does not apply to all proteins. For the antifreeze protein it can be shown that there are only minor differences between these different approaches (Fig 5; SBB-PS P value 0.0123; SCC-PS P value 0.118).

**Figure 5 a, b:**
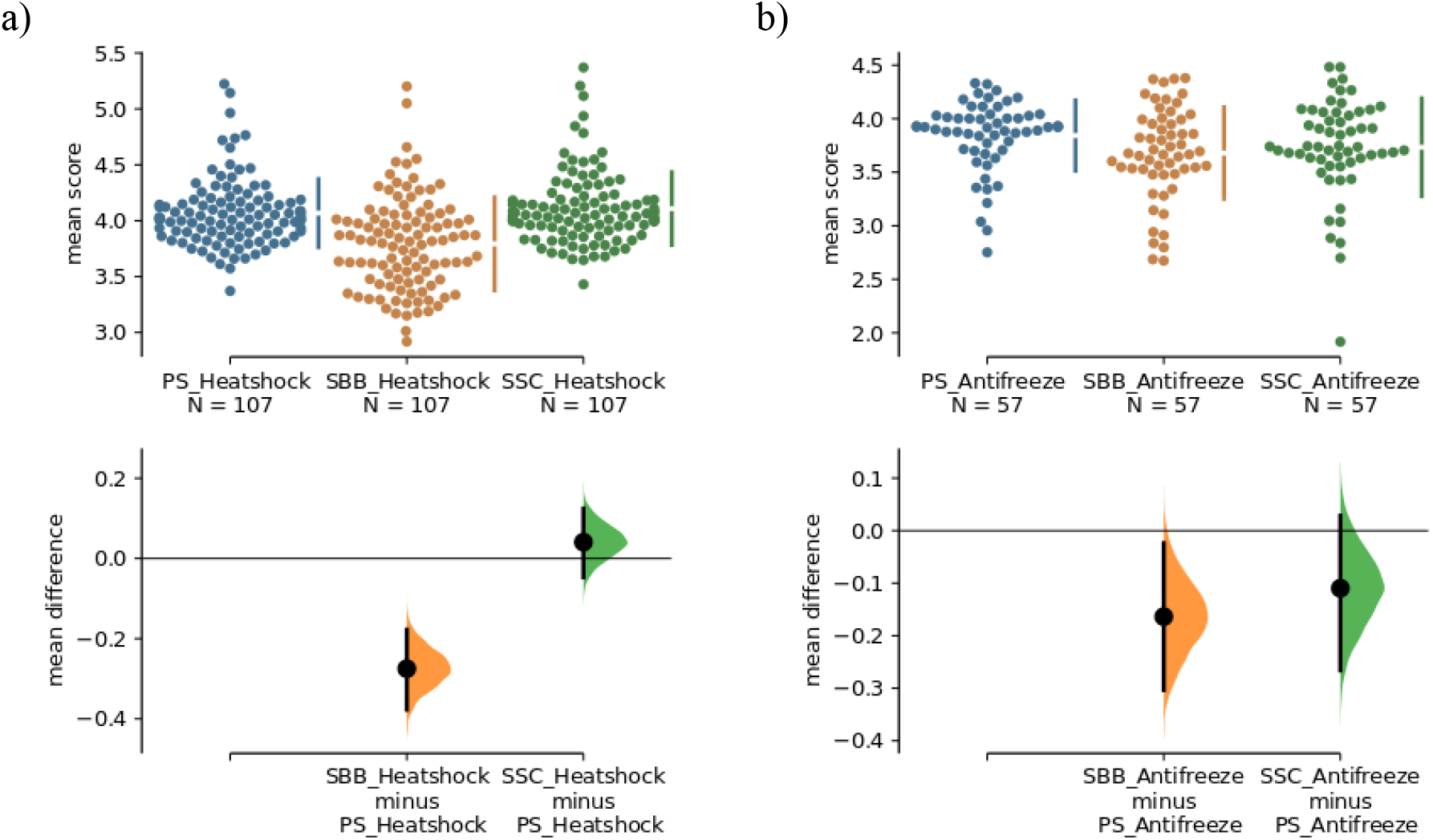
The mean difference of the surface comparisons (PS, SBB, SCC) for the heatshock and antifreeze protein are shown in the above Cumming estimation plot. The raw data is plotted on the upper axes. Each mean difference is depicted as a dot. Each 95% confidence interval is indicated by the ends of the vertical error bars. On the lower axes, mean differences are plotted as bootstrap sampling distributions (5000 bootstrap samples; confidence interval is bias-corrected and accelerated).

## Discussion

### Protein core and protein surface properties

Our hypothesis was that PS involves evolutionarily less susceptible AAs than the PC because it interacts more with the environment (Fig. 6). We could confirm this hypothesis for the mean values of the most proteins (see Results). However, there are some exceptions with regard to the sorting by organisms (see next subsection). It should be noted that the organisms (unicellular versus multicellular) may be exposed to different environments. However, there are proteins that differ from our hypothesis and for which the surface is important for protein interaction[36]. This can lead to a change in the susceptibility of the PS. This is desirable, for example, in order to subsequently industrially modify proteins[37]. An important benefit for synthetic biology is the knowledge about the susceptibility of proteins to oxidation in connection with storage and the associated degradation[16]. It is also known that proteins with susceptible AAs (tyrosine) on the surface are relevant for biological aging and age-related disease[38].

**Figure 6:**
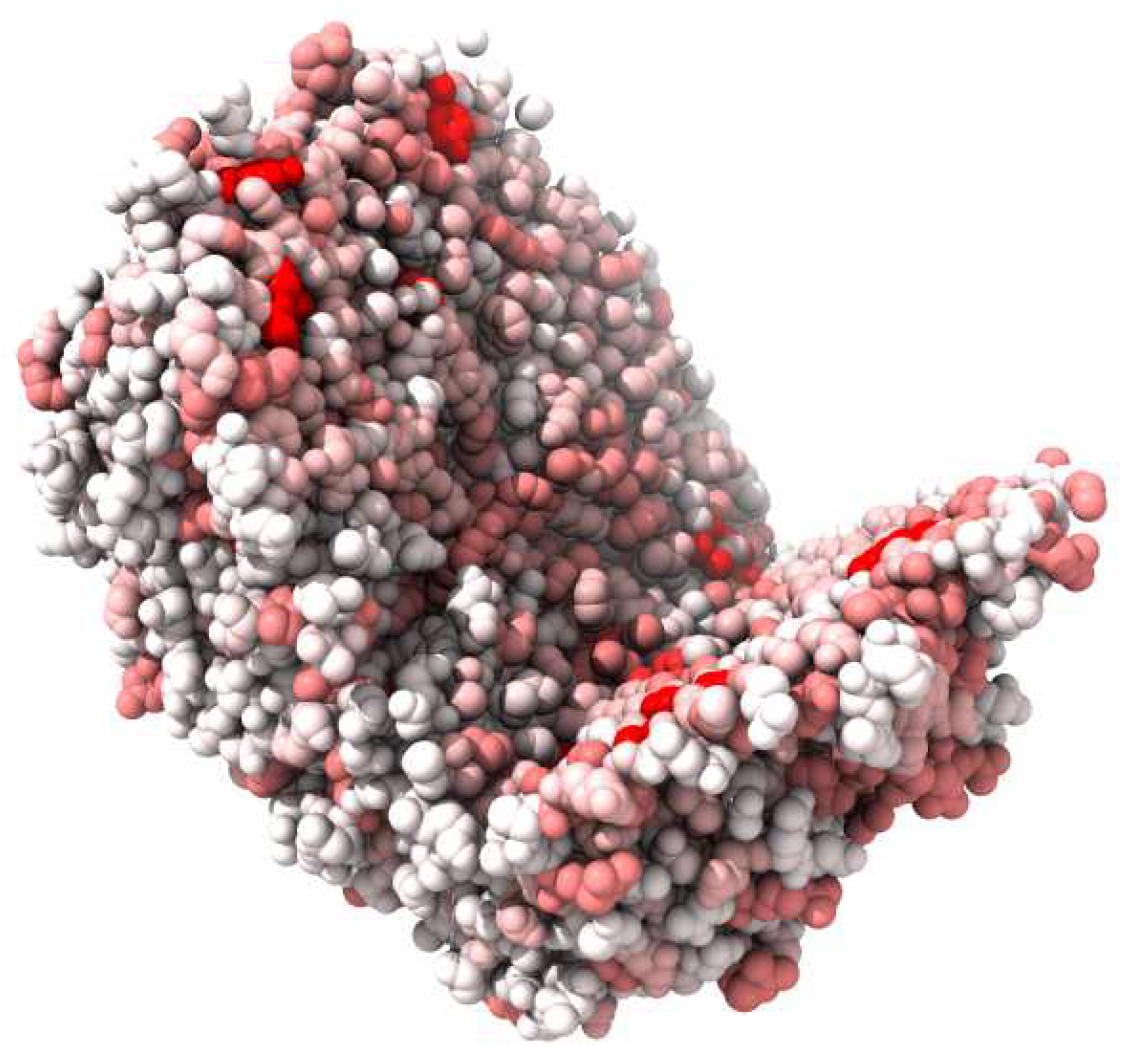
Cut of the Protein almond Pru1 (PDB entry: pdb3fz3) that has a strong diversity between the surface (low susceptibility – white) and the core (high susceptibility – red).

An important argument concerning the PS is how far it lies in or on membranes and how much it is protected by them. For example, the proteins of the respiratory chain are embedded in membranes while one side is in contact with a more aggressive environment[39]. It is well known that many proteins in the intermembrane space (IMS) contain conserved cysteine-rich sections[40]. There is still no explanation for the function of these tracts[40]. It should be noted that according to our previous results, the second most susceptible AA is cysteine [10]. Specific information on membranes could not be taken into account by our approach because the exact location of every protein was not in the data base. The same applies to complexes, which can protect parts of the protein surface from modification, except for the ones that where contained in the data base.

### Further properties – enzymes

The distinction between protein surface and core is particularly important for enzymes, since the active site is usually hidden in pockets and is either presented by a conformational change or accessible by the key-lock principle. Here no distinction is made between enzymatic and non-enzymatic proteins and thus the active sites are not examined in detail. But these hidden pockets can be an advantage in addition to substrate specificity in terms of avoiding modifications. One example is the ALDH enzyme, where the oxidation of a specific AA (Cys302) inactivates the enzyme[28]. In that paper, it was shown that neighbouring cysteine AAs protect the catalytic cysteine by covalent bonds.

When it comes to protein damage, a loss of functionality does not necessarily imply a heavy damage leading to aggregation. The required modifications for that purpose are very variable. On one hand, it was stated that nearly every modification leads to a loss of function due to changes in the conformation or in the functional domain[13]. For this, the active and passive forms would have to be taken into account. On the other hand there is an earlier finding of multiple methionine modifications not leading to a loss of function[27]. Altogether we suspect that functionality has had higher priority compared to robustness in evolution.

### Organism specific issues

The analysis, taking into account the organisms revealed a differentiation with regard to susceptibility, which could be due to the multicellularity and the adaptation to the environment (Suppl. Fig. 9). As the analysis of the PS and PC data suggested, the environment for unicellular organisms showed a shift of surface/core susceptibilities. For the individual consideration of the organisms a differentiated picture may occur. For example in *Sulfolobus* against our hypothesis the PS is more susceptible than PC in contrast to most of the other organisms (Suppl. Fig. 10). The genus *Sulfolobus* is characterized by optimal growth rate at pH 2-3 and temperatures of 70-75 °C[41]. While it keeps a pH-value of 6.5 in the cytosol[42], the high temperature could be related with the more susceptible PS. This susceptibility pattern also applies in attenuated form to all other unicellular organisms which, in contrast to multicellular organisms, have a higher reproduction rate. This allows them to get rid of damaged proteins by asymmetric division or apoptosis. Multicellular organisms, on the other hand, do the same, but the damaged proteins remain in the intercellular spaces or have to be expensively transported away (immune cells, [43,44]). We hypothetize that the outer cell layers (e.g. a major constituent of the skin is collagen) of multicellular organisms have a typical proteome that protects them from an aggressive environment.

### Possible extensions

In this paper we have investigated the susceptibility of AAs in proteins with respect to their spatial location. A further interesting classification model is quantitative structure-activity relation (QSAR) analysis[45,46]. Here, in contrast to our analysis, a relationship is established between the structure and the activity. This can also be combined with our classification model relating structure and aging (susceptibility).

Another approach would be to describe the exact spatial orientation of the AA susceptibilities using a tensor. The different proteins could be sorted according to linear, planar or spherical spatial susceptibility. It is to be expected that e.g. membrane proteins (Fig. 7), which pass through the membrane, could show a linear part and surface-associated proteins show rather planar properties. Unbound proteins are more likely to have spherical susceptibilities because they can be attacked from all sides.

**Figure 7:**
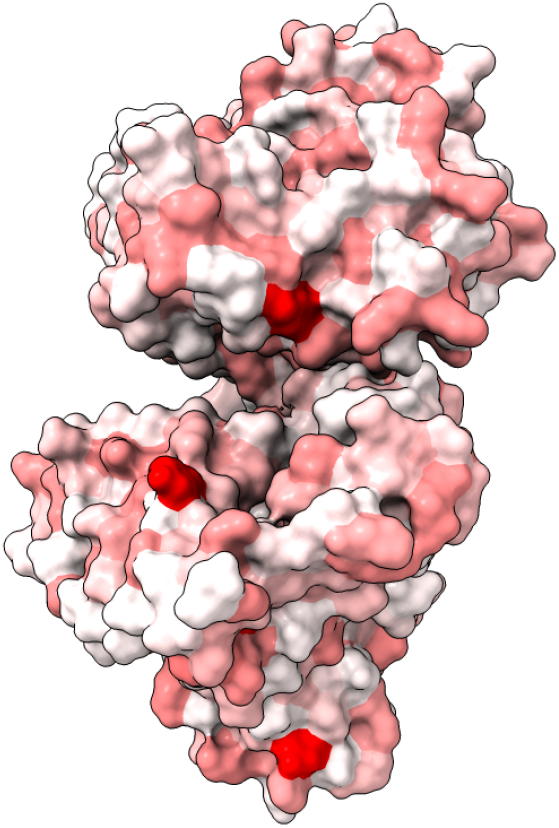
Surface susceptibility of the protein phosphodiesterase 4B (PDB entry: pdb5ohj)

A further approach would be to study the accessibility on the atomic level in the form of a spherical representation with calculated surface fraction[22]. This is a possible extension of our work since the surface fractions can be used as weightings and the single atoms can be scored in terms of their susceptibility. For reasons of simplification, however, we have limited ourselves to the AAs. This has the advantage that the inaccuracy of the spatial structure has less influence on the weighting. Inaccuracies can be e.g. the degrees of freedom of individual AAs, as well as the free energy, the volume or the entropy. An example are the transcription factors (some of them involve many random coils), which often have an undefined surface and are therefore difficult to calculate.

## Acknowledgements

We thank all the colleagues of the department of bioinformatics. Also we like to thank Steve Hoffmann, Lars-Oliver Klotz, Holger Steinbrenner and André Then for helpful discussions.

## List of abbreviations

3D: three-dimensional
AA: amino acid
PC: protein core
PDB: protein data bank
PS: protein surface
SBB: surface backbone
SD: standard deviation
SSC: surface side chain
WP: whole protein

