## Supplemental information for "Influence of spatial structure on protein damage susceptibility – A bioinformatics approach"

Table of Contents Page

1. Data preparation S2

2. Hydrophobicity classification S4

3. Further amino acid distribution visualizations S5

4. Further comparison of protein surface and core properties S7

5. Further comparison of enzyme properties S8

6. Comparison between organisms S9

7. Appendix S10

8. References S17

#### 1. Data preparation

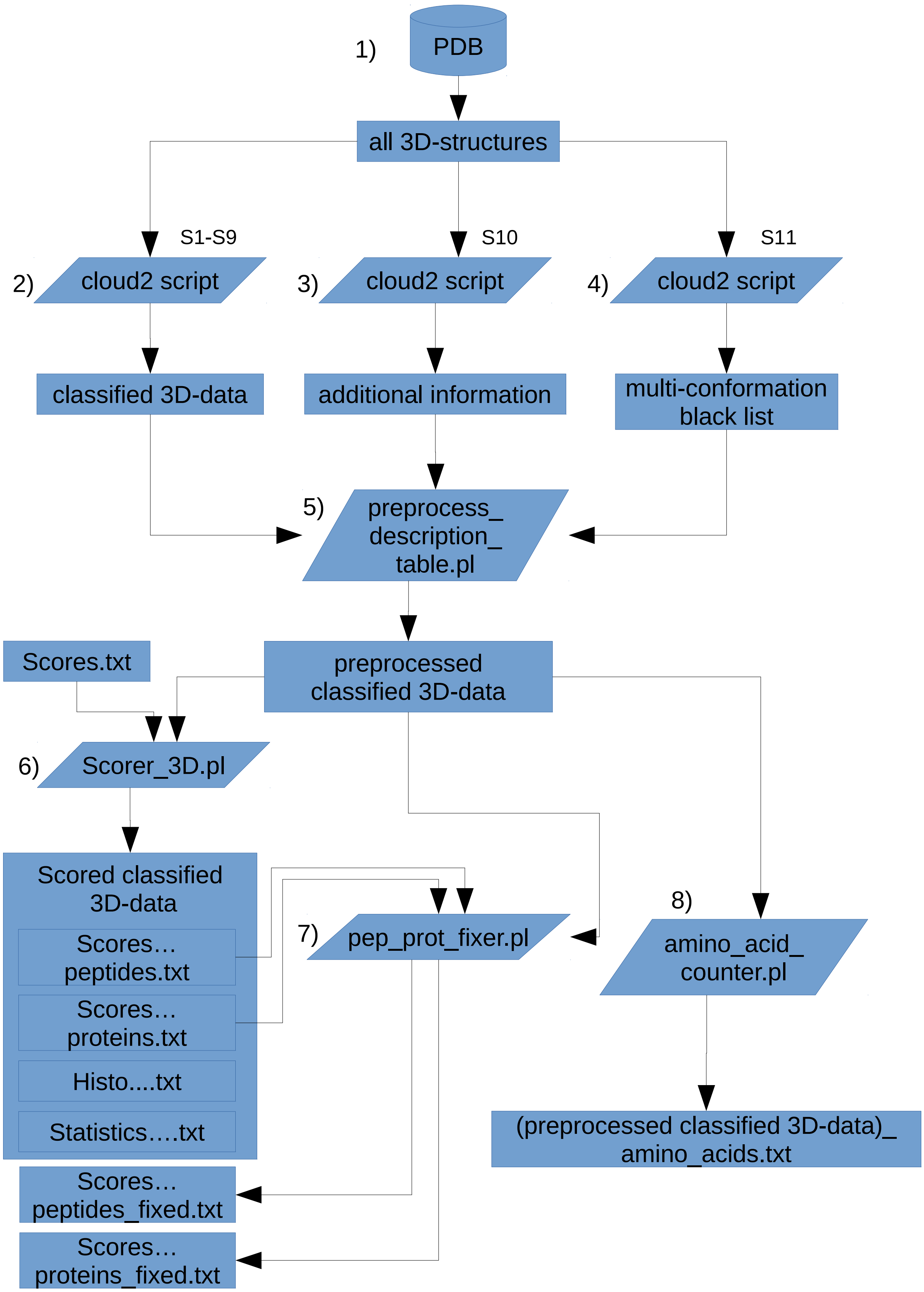

**Suppl. Figure 1:** Flowchart of the whole process of generating the scores and counting of the amino acids.

1) The spatial data was downloaded from the Protein Data Bank (PDB) (ftp.wwpdb.org/pub/pdb/data/structures/all/pdb; 02.05.2018).

2) The scripts #1 – #9 (see Appendix) were used in the program Cloud2 Version 14.3.20 (Heiko Stark, Jena, Germany, URL: https://starkrats.de) to extract the respective amino acids (AA). This resulted in the different sequence files ('all', 'core', 'hull', ...) modified with the respective spatial information. This leads to 9 different data sets, but for simplicity it is summarized here as classified 3D-data. That means the following steps have to be done separately for every data set.

3)The script #10 was used to retrieve additional information like descriptions and organisms out of the structure files to include them later.

4) Multi-conformational entries could not be handled at this point, that is why script #11 was used to make a list of all multi-conformational IDs to exclude these sequences later.

Step #2 to #4 can also be done in one, but due to the development process we came to this modularised approach which gives the possibility to alter one aspect without having to recalculate all the spatial data.

5) The perl (v5.22.1) program 'preprocess_description_table.pl' combines the gathered information from step #2 to #4. It adds the additional information to the headers of the calculated sequences and removes the sequences with the IDs from the multi-conformation black list. The resulting files now have the ending '..._descr...' added to them but are here listed as preprocessed classified 3D-data.

6) The program 'Scorer_3D.pl' is a modification of the scoring program from Fichtner et al. (2019) [1]. It leaves out the GO-annotation. It also uses the scores from the table in 'Scores.txt'. For every data set a peptide and protein score file is created as well as a histogram and a statistics file. See also supplement of Fichtner et al. (2019)[1] for further details.

7) Because the sequences are reduced in step #2 (except for the data set 'all') the allocation to peptides and proteins is not working as intended in step #6. That means proteins can be degraded to peptides if they fall under the threshold of 100 AAs. The program 'pep_prot_fixer.pl' is there to reallocate the false positive peptides. It takes the peptide and the protein score file, compares them with the preprocessed classified 3D-data and does the necessary arrangements. It also removes all entries with an 'X'-amount higher than the chosen threshold. In our case we decided to use 0.05 as a default value. The created score files have the ending '..._fixed...' and are the final result in the chain.

8) An additional consideration is made in step #8 out of the 'preprocessed classified 3D-data'. The program 'amino_acid_counter.pl' is counting the occurrence of all characters. The results are discussed in our paper and in Suppl. 3 there are additional diagrams for this approach.

#### 2. Hydrophobicity classification

We try to gather information about the amount of hydrophobic AAs for every protein to give a relative representation of the bonding potential of hydrophobic interactions that lie within the proteins. When it comes to conformational changes this can lead very easily to aggregations. Beware this information is not oriented on protein surface hydrophobicity! It is generally known that there is a difference in the distribution of hydrophobic versus hydrophilic AAs[2]. This is because proteins have an aqueous environment and therefore have to be hydrophilic.

Concerning hydrophobicity there is no definite classification. Here we are holding on to the description of Voet and Voet[3], Lesk[4] and Berg, Stryer and Tymoczko[5] with some minor adjustments because we are only interested in hydrophobic potential. That means all the nonpolar AAs which are alanine, phenylalanine, glycine, isoleucine, leucine, methionine, proline and valine are definitely classified as hydrophobic. Two other borderline cases namely tyrosine and tryptophan are also seen as hydrophobic due to their aromatic rings (Tab. 1).

Glycine for example has a side chain which is so small that the attributes of the backbone dominate. That means in unbound form it would be hydrophilic and when bound it would be hydrophobic. Since here we are working with bound AAs the classification is decided to be hydrophobic. A contradiction for the first and last AA in the chain will be neglected.

**Suppl. Table 1:** The upper table shows the properties of the amino acids with regard to hydrophobicity. The lower table shows the classifications of amino acids with regard to hydrophobicity in the literature. The representation of the bold entries show the own decisions about different views in the literature regarding hydrophobicity.

| amino acid | A | C | D | E | F | G | H | I | K | L | M | N | O | P | Q | R | S | T | U | V | W | Y |
|---|---|---|---|---|---|---|---|---|---|---|---|---|---|---|---|---|---|---|---|---|---|---|
| hydrophobic | X | | | | X | X | | X | | X | X | | | X | | | | | | X | X | X |
| hydrophilic | | X | X | X | | | X | | X | | | X | | | X | X | X | X | | | | X |

| amino acid | A | C | D | E | F | G | H | I | K | L | M | N | O | P | Q | R | S | T | U | V | W | Y |
|---|---|---|---|---|---|---|---|---|---|---|---|---|---|---|---|---|---|---|---|---|---|---|
| Voet & Voet | X | | | | X | X | | X | | X | X | | | X | | | | | | X | X | |
| Lesk | X | | | | X | X | | X | | X | X | | | X | | | | | | X | | |
| Stryer | X | | | | X | X | | X | | X | X | | | X | | | | | | X | X | |
| This work | X | | | | X | X | | X | | X | X | | | X | | | | | | X | **X** | **X** |

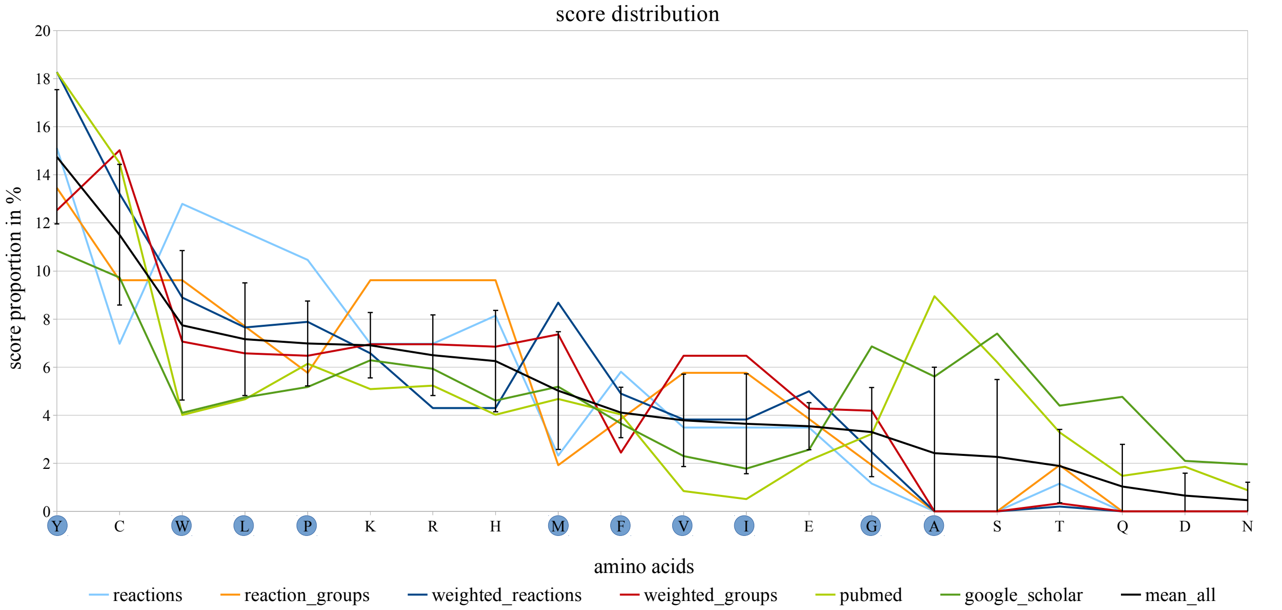


**Suppl. Figure 2:** Score distribution from Fichtner et al. (2019) sorted by score. The amino acids classified by us as hydrophobic are marked with blue circles.

#### 3. Further amino acid distribution visualizations

In the PDB data the information is resolved on the level of atoms which are allocated to different AAs. There are also other entries than the 20 standard AAs listed. This includes next to the very rare selenocysteine ('U'; only 191 occurrences) and pyrrolysine ('O', only 2 occurrences) also unknown positions ('X'). Even other molecules like ligands or water on the protein surface are listed. These entries are put together in one 'X' at the end of the sequence by cloud2. For the creation of the diagrams only the information for the 20 standard AAs was considered. That means especially in the calculation 'X's for example were not considered.

For a closer look at AA distributions Suppl. Fig. 3 shows the distribution of all standard AAs over all proteins we analyzed from PDB.

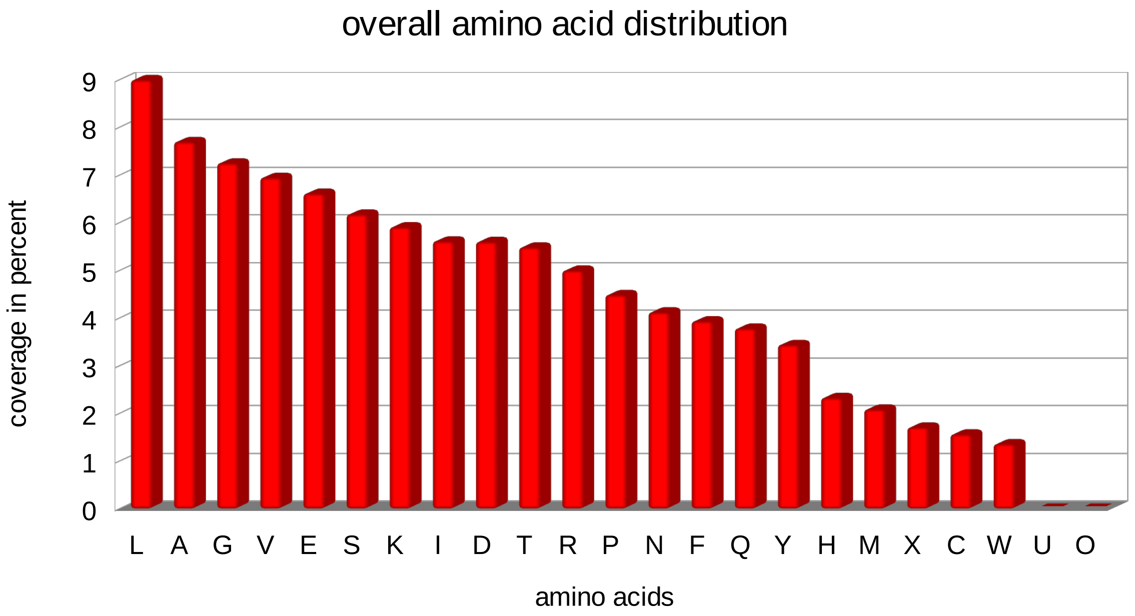


**Suppl. Figure 3:** Normalized amino acid distribution for all proteins with structure information from PDB.

In the following some normalized versions for the AA distributions with different focuses are shown. Suppl. Fig. 4 shows how the AAs distribute over protein surface (PS) and core (PC). Here a normalization is shown where the real values of an AA for PS and PC are divided by the sum of all values for that AA. In addition, it was changed into percent where PS+PC for every AAs equals 100 %. This shows the distribution of the respective AA over PS and PC. As expected the PC dominates most of the entries which is because the complete data set of PDB contains many bigger proteins. With increasing size the share of AAs considered to be in the core increases in general much more than in the PS which leads to the PC containing most of the AAs. That is why another normalization step was made.

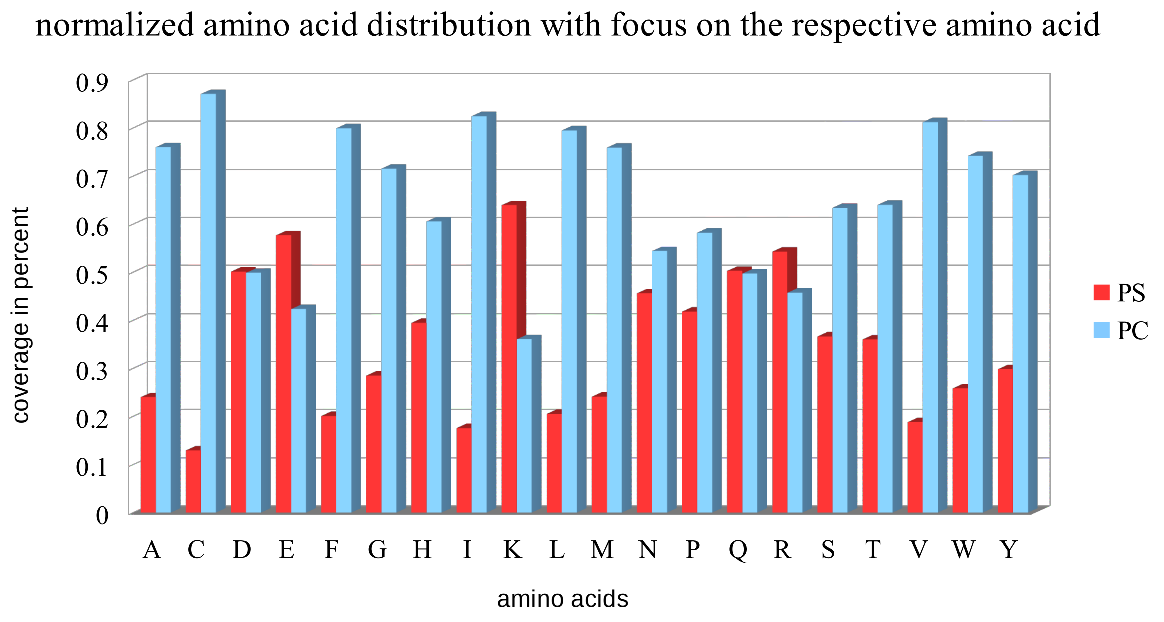


**Suppl. Figure 4:** Normalized amino acid distribution with focus on the respective amino acid. The protein surface (PS) and core (PC) value for every amino acid add up to 100 %. For example for all alanine over all proteins 24 % distribute on the PS and 76 % on the PC.

The next figure (Suppl. Fig. 5) focuses on the other direction. Here, it is shown how the PS and PC distribute over all the AAs. Therefore, the real values of an AA for PS and PC are divided by the sum of all respective PS or PC values. In addition, it was changed into percent where all the values for PS/PC together equal 100 %. This shows the distribution of the surface or the core over all AAs. This normalization is the best to compare the surface with the PC proportionally.

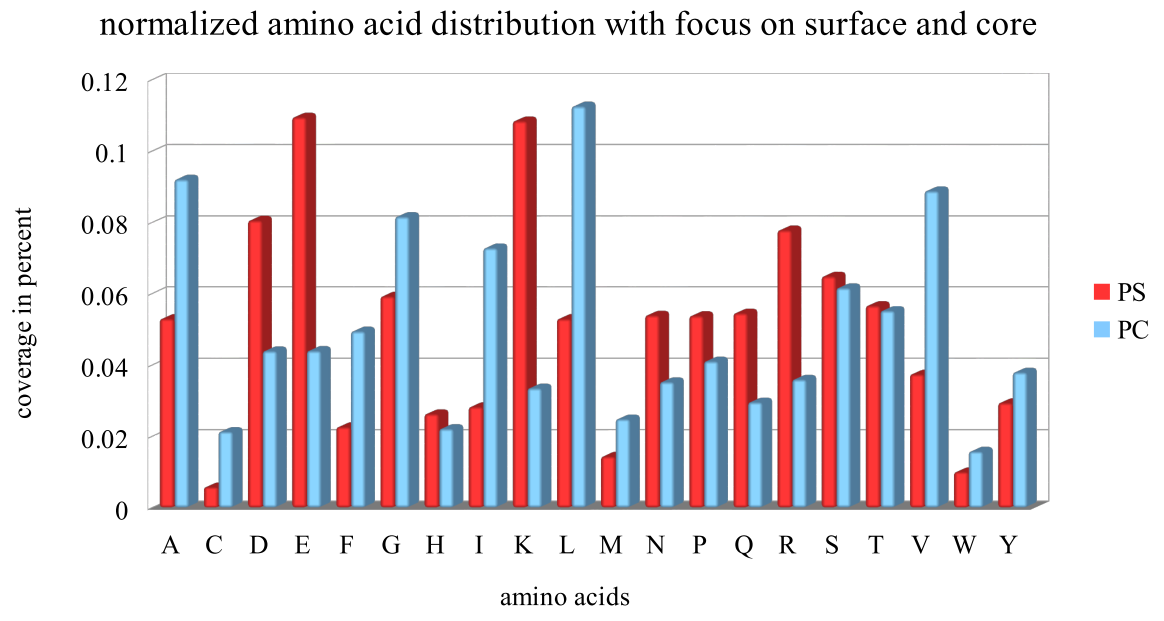


**Suppl. Figure 5:** Normalized amino acid distribution with focus on protein surface (PS) and core (PC). All red columns together add up to 100 % (PS) and all blue columns together add up to 100 %

(PC). For example for all amino acids on the PS about 5 % are alanine and of all amino acids in the PC about 9 % are alanine.

#### 4. Further comparison of protein surface and core properties

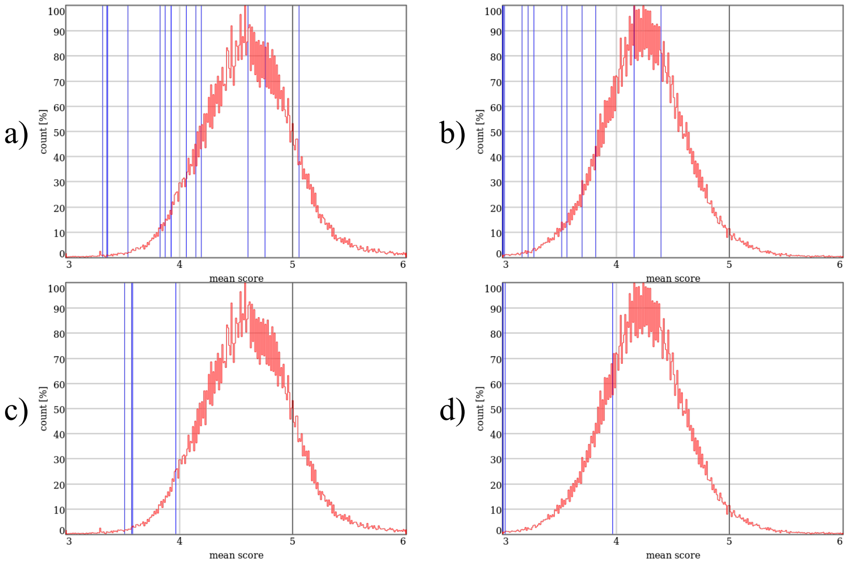


**Suppl. Figure 6:** Diagrams of the mean score of protein core (PC) and protein surface (PS) of flaggellin and spidroin. a) PC flagellin, b) PS flagellin, c) PC spidroin, d) PS spidroin. The red graph shows the histogram of all proteins and the vertical blue lines indicate the position of the protein entries. The black line indicates the mean score of a random artificial protein.

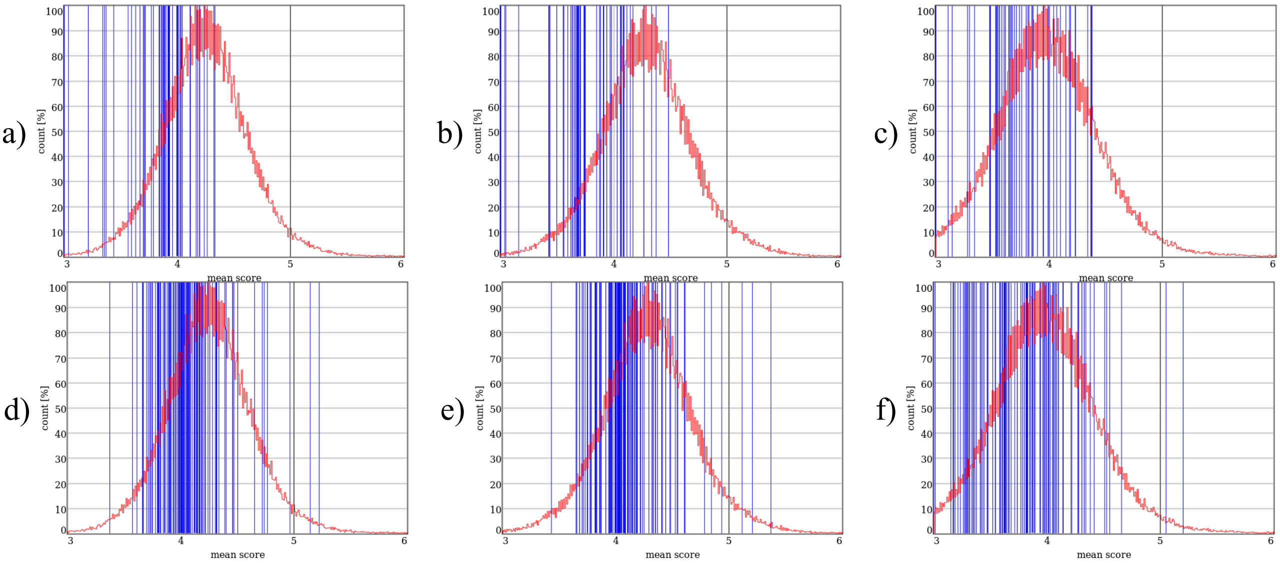


**Suppl. Figure 7:** Diagrams of the mean score of surface types protein surface (PS), surface backbone (SBB), surface side chain (SSC) for antifreeze and heat shock. a) PS antifreeze, b) SBB antifreeze, c) SSC antifreeze, d) PS heat shock e) SBB heat shock f) SSC heat shock. The red graph shows the histogram of all proteins and the vertical blue lines indicate the position of the protein entries. The black line indicates the mean score of a random artificial protein.

### 5. Further comparison of enzyme properties

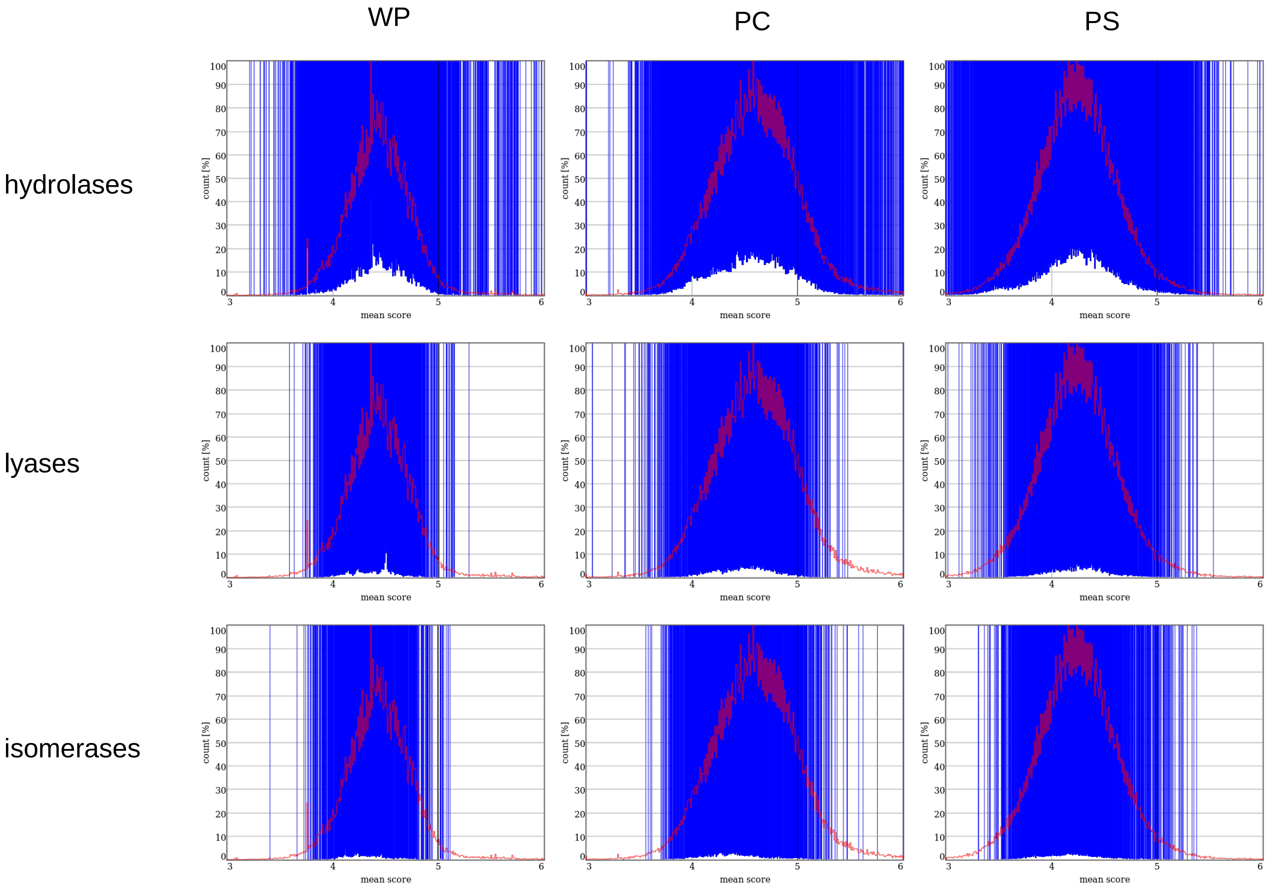


**Suppl. Figure 8:** Diagrams of the mean score of whole protein (WP), protein core (PC) and protein surface (PS) of hydrolases (EC:03), lyases (EC:04) and isomerases (EC:05). The red graph shows the histogram of all proteins and the vertical blue lines indicate the position of the protein entries. The black line indicates the mean score of a random artificial protein.

For more examples we refer to our website http://damage.stark-jena.de.

#### 6. Comparison between organisms

Our analysis revealed visible differences between organisms from different kingdoms (Suppl. Fig. 9). Bacteria show more unsusceptible proteins than other organisms, while animals show more susceptible proteins. The plants, which seem neither susceptible nor unsusceptible, occupy a special position.

*Calculation and normalization*

The calculation for the Supplement Figures 9 and 10 were calculated in two cases. The two cases take into account the difference between the relation of the mean score for the protein surface (PS) and the protein core (PC). In case 2 a normalization has been made, to show all values in the range of 1 to 2 and with that overall in the range of 0 to 2.

case 1 mean score PS < mean score PC : mean score PS / mean score PC
case 2 mean score PS > mean score PC : 1+(1- (mean score PC / mean score PS))

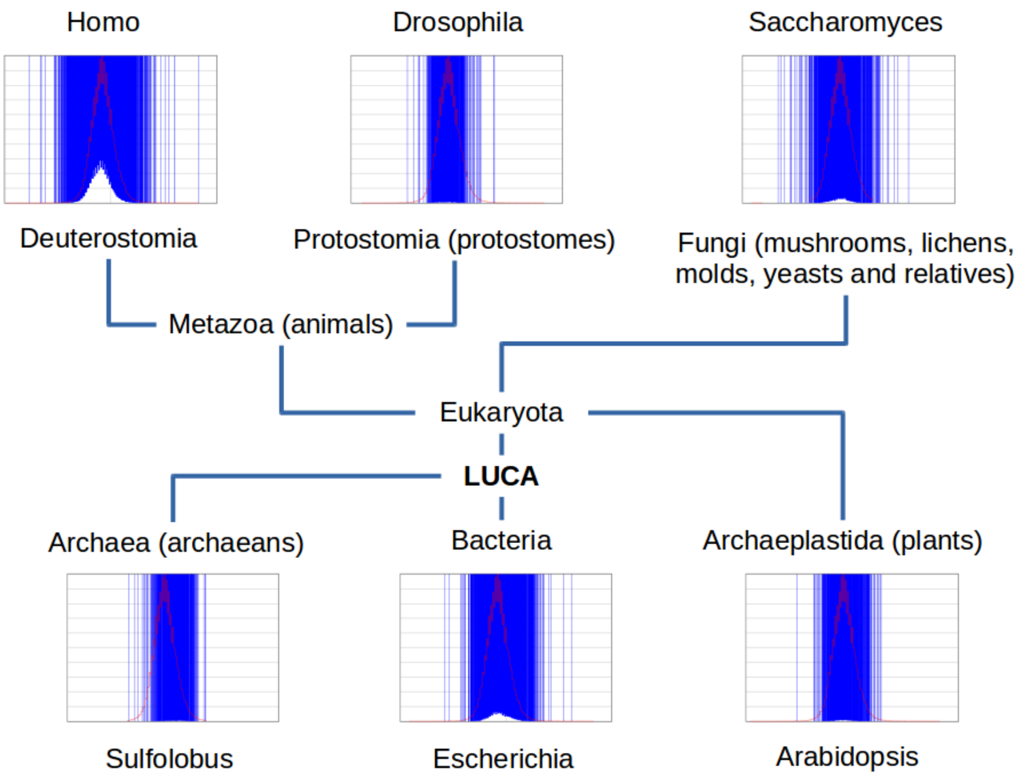


**Suppl. Figure 9:** Cladogram of the genera *Sulfolobus, Escherichia, Arabidopsis, Saccharomyces, Drosophila and Homo*. The leafs/clades show the distribution of the peptides and proteins with regard to the difference between protein core (PC) and surface (PS) (see difference calculation).

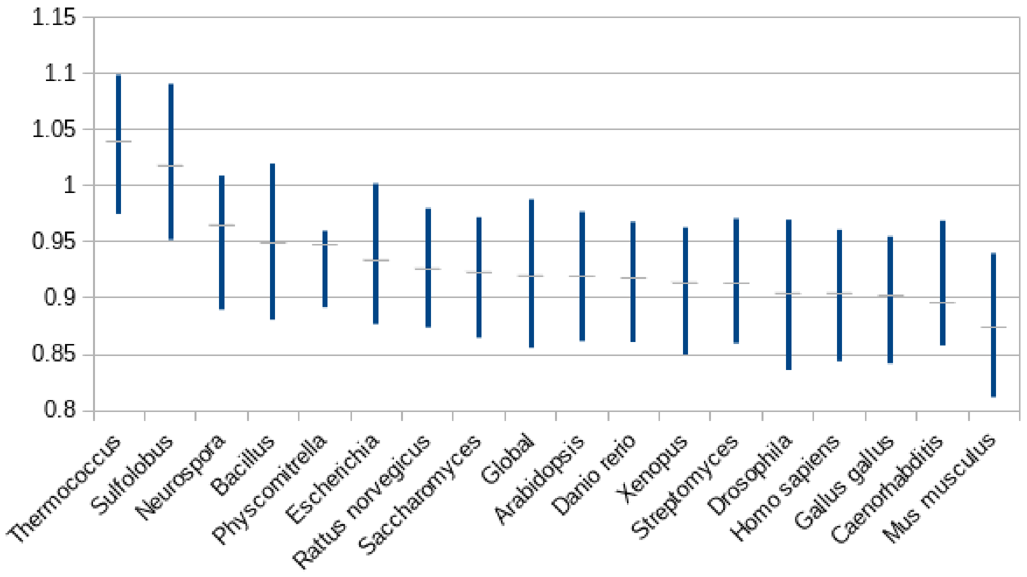


**Suppl. Figure 10:** Sorted representation of the organisms with regard to the susceptibility between protein surface (PS) and protein core (PC) (see difference calculation). Values above 1 mean a more susceptible PS compared to the PC and values below 1 mean a more susceptible PC compared to the PS.

A more detailed analysis of the distribution shows differences between organisms in the susceptibilities in PS and PC. These clearly occur in the *Archea*, since on average they have a more susceptible PS than the PC (Suppl. Fig. 10). It is noticeable that *Rattus norvegicus* and *Mus musculus* differ greatly from their closest relatives (this may be due to the fact that these are experimental animals and therefore certain protein classes are overrepresented). Otherwise, the organisms are divided into unicellular and multicellular groups.

#### 7. Appendix

**Script 1:** all

```
io.files *.ent.gz
io.files.sort

for io.files.length

   new.all
   set.red
   filename := io.workdir / io.files.getdir / io.files.getfile
   import.pdb.atom filename

   filename := io.workdir / io.files.getdir / 'result_all.txt'
   append.aminoacids filename

   io.files.next

end
```

**Script 2:** core_6

```
io.files *.ent.gz
io.files.sort

for io.files.length

        new.all
        set.red
        filename := io.workdir / io.files.getdir / io.files.getfile
        import.pdb.atom filename
        edit.name.to.unique
        set.ref.green
        bool.copy.layer

        set.ref.purple
        mesh.point.hull.x <0, 0, 0> 3 6 0
        mesh.point.hull.y <0, 0, 0> 3 6 0
        mesh.point.hull.z <0, 0, 0> 3 6 0
        set.purple
        edit.erase.duplicates

    // mask aminoacids
        set.red
        set.ref.purple
        bool.inside.circle
        bool.mask.points

    // mask core aminoacids
        set.green
        set.ref.red
        bool.outside.circle
        bool.mask.objects

        filename := io.workdir / io.files.getdir / 'result_core_6.txt'
        append.aminoacids filename

        io.files.next

    end
```

**Script 3:** core_7

```
    io.files *.ent.gz
    io.files.sort

    for io.files.length

        new.all
        set.red
        filename := io.workdir / io.files.getdir / io.files.getfile
        import.pdb.atom filename
        edit.name.to.unique
        set.ref.green
        bool.copy.layer

        set.ref.purple
        mesh.point.hull.x <0, 0, 0> 3.5 7 0
        mesh.point.hull.y <0, 0, 0> 3.5 7 0
        mesh.point.hull.z <0, 0, 0> 3.5 7 0
```

```
    set.purple
    edit.erase.duplicates

// mask aminoacids
    set.red
    set.ref.purple
    bool.inside.circle
    bool.mask.points

// mask core aminoacids
    set.green
    set.ref.red
    bool.outside.circle
    bool.mask.objects

    filename := io.workdir / io.files.getdir / 'result_core_7.txt'
    append.aminoacids filename

    io.files.next

end
```

**Script 4:** hull_6

```
io.files *.ent.gz
io.files.sort

for io.files.length

    new.all
    set.red
    filename := io.workdir / io.files.getdir / io.files.getfile
    import.pdb.atom filename

    set.ref.purple
    mesh.point.hull.x <0, 0, 0> 3 6 0
    mesh.point.hull.y <0, 0, 0> 3 6 0
    mesh.point.hull.z <0, 0, 0> 3 6 0
    set.purple
    edit.erase.duplicates
    set.red
    set.ref.purple
    bool.inside.circle
    bool.mask.points

    filename := io.workdir / io.files.getdir / 'result_hull_6.txt'
    append.aminoacids filename

    io.files.next

end
```

**Script 5:** hull_7

```
io.files *.ent.gz
io.files.sort

for io.files.length
```

```
    new.all
    set.red
    filename := io.workdir / io.files.getdir / io.files.getfile
    import.pdb.atom filename

    set.ref.purple
    mesh.point.hull.x <0, 0, 0> 3.5 7 0
    mesh.point.hull.y <0, 0, 0> 3.5 7 0
    mesh.point.hull.z <0, 0, 0> 3.5 7 0
    set.purple
    edit.erase.duplicates
    set.red
    set.ref.purple
    bool.inside.circle
    bool.mask.points

    filename := io.workdir / io.files.getdir / 'result_hull_7.txt'
    append.aminoacids filename

    io.files.next

end
```

**Script 6:** hull-backbone_6

```
io.files *.ent.gz
io.files.sort

for io.files.length

    new.all
    set.red
    filename := io.workdir / io.files.getdir / io.files.getfile
    import.pdb.backbone filename

    set.ref.purple
    mesh.point.hull.x <0, 0, 0> 3 6 0
    mesh.point.hull.y <0, 0, 0> 3 6 0
    mesh.point.hull.z <0, 0, 0> 3 6 0
    set.purple
    edit.erase.duplicates
    set.red
    set.ref.purple
    bool.inside.circle
    bool.mask.points
    bool.erase.line.materials -100 0

    filename := io.workdir / io.files.getdir / 'result_hull-backbone_6.txt'
    append.aminoacids filename

    io.files.next

end
```

**Script 7:** hull-backbone_7

```
io.files *.ent.gz
io.files.sort

for io.files.length

   new.all
   set.red
   filename := io.workdir / io.files.getdir / io.files.getfile
   import.pdb.backbone filename

   set.ref.purple
   mesh.point.hull.x <0, 0, 0> 3.5 7 0
   mesh.point.hull.y <0, 0, 0> 3.5 7 0
   mesh.point.hull.z <0, 0, 0> 3.5 7 0
   set.purple
   edit.erase.duplicates
   set.red
   set.ref.purple
   bool.inside.circle
   bool.mask.points
   bool.erase.line.materials -100 0

   filename := io.workdir / io.files.getdir / 'result_hull-backbone_7.txt'
   append.aminoacids filename

   io.files.next

end
```

**Script 8:** hull-side_chain_6

```
io.files *.ent.gz
io.files.sort

for io.files.length

   new.all
   set.red
   filename := io.workdir / io.files.getdir / io.files.getfile
   import.pdb.backbone filename

   set.ref.purple
   mesh.point.hull.x <0, 0, 0> 3 6 0
   mesh.point.hull.y <0, 0, 0> 3 6 0
   mesh.point.hull.z <0, 0, 0> 3 6 0
   set.purple
   edit.erase.duplicates
   set.red
   set.ref.purple
   bool.inside.circle
   bool.mask.points
   bool.erase.line.materials 0 100

   filename := io.workdir / io.files.getdir / 'result_hull-side_chain_6.txt'
   append.aminoacids filename

   io.files.next
```

```
end
```

**Script 9:** hull-side_chain_7

```
io.files *.ent.gz
io.files.sort

for io.files.length

   new.all
   set.red
   filename := io.workdir / io.files.getdir / io.files.getfile
   import.pdb.backbone filename

   set.ref.purple
   mesh.point.hull.x <0, 0, 0> 3.5 7 0
   mesh.point.hull.y <0, 0, 0> 3.5 7 0
   mesh.point.hull.z <0, 0, 0> 3.5 7 0
   set.purple
   edit.erase.duplicates
   set.red
   set.ref.purple
   bool.inside.circle
   bool.mask.points
   bool.erase.line.materials 0 100

   filename := io.workdir / io.files.getdir / 'result_hull-side_chain_7.txt'
   append.aminoacids filename

   io.files.next

end
```

**Script 10:** table

```
io.files *.ent.gz
io.files.sort

tab := '          '

store '../table.pdb'
@ 'ID' tab '3DID' tab 'EC' tab 'CLASS' tab 'TITLE' tab 'ORGANISM'
end

for io.files.length

   new.all
   filename := io.workdir / io.files.getdir / io.files.getfile
   import.pdb.atom filename

   filename := '../table.pdb'
   append filename
@ io.files.getname tab pdb.id tab pdb.ec tab pdb.class tab pdb.title tab pdb.organism
   end

   io.files.next

end
```

**Script 11:** multimodels

```
io.dirs.complete * // load all sub dirs into a dir list

io.files *.ent.gz

output.store "models.txt"

for io.files.length // for all files
	filename := io.workdir / io.files.getdir / io.files.getfile
	bool := text.find filename "MODEL        2" "MODEL        3"
	if bool
	then print filename
	io.files.next
end

end
```
